# RNA-mediated nucleosome depletion is required for elimination of transposon-derived DNA

**DOI:** 10.1101/2022.01.04.474918

**Authors:** Aditi Singh, Xyrus X. Maurer-Alcalá, Therese Solberg, Silvan Gisler, Michael Ignarski, Estienne C. Swart, Mariusz Nowacki

## Abstract

Small RNAs are known to mediate silencing of transposable elements and other genomic loci, increasing nucleosome density and preventing undesirable gene expression. Post-zygotic development of the *Paramecium* somatic genome requires elimination of thousands of transposon remnants (IESs) and transposable elements that are scattered throughout the germline genome (Garnier et al. 2004). The elimination process is guided by Piwi-associated small RNAs and leads to precise cleavage at IES boundaries (Bouhouche et al. 2011; Furrer et al. 2017). Previous research suggests that small RNAs induce heterochromatin formation within IESs, which, in turn, is required for DNA elimination (Liu et al. 2007). Here we show that IES recognition and precise excision is facilitated by recruitment of a homolog of a chromatin remodeler ISWI, which depletes target genomic regions of nucleosomes, making the chromatin accessible for DNA cleavage. ISWI knockdown in *Paramecium* leads to pronounced inhibition of DNA elimination. Furthermore, nucleosome profiling indicates that ISWI is required for IES elimination in nucleosome-dense genomic regions, while other IESs do not require small RNAs or ISWI for excision. ISWI silencing notably also reduces DNA elimination precision, resulting in aberrant excision at alternative IES boundaries. In summary, we demonstrate that chromatin remodeling that increases DNA accessibility together with small RNAs are necessary for efficient and precise DNA elimination in *Paramecium*.

## Introduction

ISWI proteins form different complexes interacting with several conserved domains, with each complex modulating a discrete function (Dirscherl and Krebs 2004).

Although ISWI complexes have distinct functions, the generalized mechanism underlying their various roles is based on altering nucleosome spacing. This moving around of the nucleosome helps DNA-binding proteins access sites that were previously unavailable (Clapier and Cairns 2009). However, the understanding of why ISWI protein complexes are chosen over other chromatin complexes or how they determine which regulatory process to target is still not clear. The study of ISWI complexes in diverse species will help to generalize the understanding of their mechanisms.

Ciliates, such as *Paramecium tetraurelia*, provide an excellent model system to understand the dynamic genome organization in eukaryotic cells due to their unique feature nuclear dimorphism. The formation of Paramecium’s somatic nucleus during sexual reproduction involves DNA endoreplication, DNA elimination, DNA repair and transcription of genes that are specifically expressed when these processes occur (Chalker and Yao 2011). Hence, the chromatin needs to be in a tightly controlled dynamic state. The germline micronuclear (MIC) genome contains regions that are removed during the development of the somatic macronuclear (MAC) genome (Beisson et al. 2010c), in a sophisticated process of genome reorganization, a natural form of genome editing. During this event, about 45,000 unique, non-coding Internal Eliminated Sequences (IES) are typically precisely excised (Arnaiz et al. 2012). Precise elimination of IESs is crucial for the formation of a functional somatic genome, since these sequences would otherwise frequently interrupt exonic coding sequences.

DNA elimination process, carried out by a catalytically active domesticated transposase, PiggyMac (*PGM) (Baudry et al. 2009)* , requires the accessibility to the boundaries of IESs. One way to make the DNA accessible for cleavage would be through the action of ATP-dependent remodelers, such as ISWI proteins, that can restructure the chromatin. Indeed, global changes of nucleosome density can act as a regulatory factor controlling the access to DNA (Sadeh and Allis 2011; Rando and Winston 2012). Therefore, one can imagine a similar mechanism existing in ciliates, which modulates nucleosome density changes and thereby transiently makes IESs accessible for excision. To this end, we studied the putative role of *Paramecium* ISWI during genome reorganisation.

*Paramecium tetraurelia* has five putative ISWI homologs with the characteristic SWI/SNF family ATPase core domain as well as SANT and SLIDE domains towards their C-termini (Figure 1A). Out of these four are pairs of paralogs arising from the well-characterized whole genome duplication (WGD, Figure 1B) events in *Paramecium (Aury et al. 2006)*. Of the paralogs, the homolog characterized here, *ISWI1*, shows substantial differential upregulation during the macronuclear development whereas *ISWI2, ISWI3* and *ISWI4* do not (Figure 1C). The remaining *ISWI* homolog, *ISWI5,* also shows substantial differential expression, peaking during meiosis and fragmentation of the parental MAC before decreasing in abundance for the remainder of development (Figure 1C).

**Figure 1.**
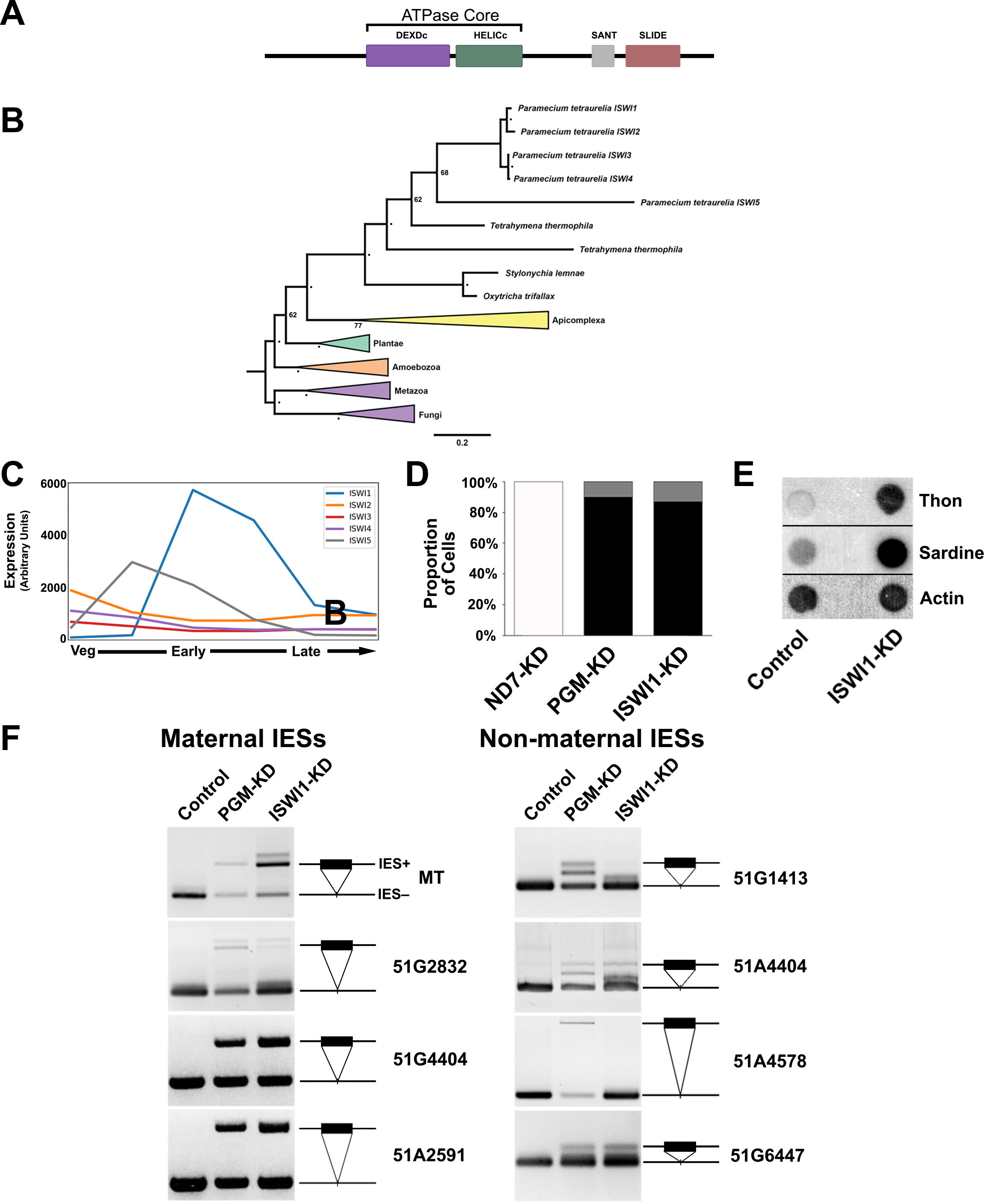
Properties of ISWI1 and ISWI1-KD effects on DNA elimination. (A) Predicted protein domains in ISWI1. (B) Phylogenetic analysis of ISWI proteins in selected organisms. Node bootstrap values below ≥ 80 are indicated by ‘•’ or are otherwise labelled. (C) Gene expression profile (in arbitrary units) of ISWI genes based on published RNA-seq data (Arnaiz et al., 2017). Veg: cells undergoing vegetative division; Early: ∼50% of cells with fragmented parental macronucleus (our early time point); Late: the majority of cells with a visible anlagen (our late timepoint). (D) Survival test graph. Dead cells are represented in black, sick in grey and normally dividing cells in white. PGM-KD is a positive control and ND7-KD is a negative control. (E) Dot blot analysis to check the effect of ISWI1-KD on transposon elimination. Probes against transposons Sardine and Thon were used while a probe against Actin was used as a loading control. (F) IES retention PCR (cropped inverted images). Four maternally-controlled IES and four non-maternally controlled IESs are shown. The IES+ band represents retained IES; the IES- band represents an excised IES; additional bands are likely PCR artefacts.

## Results

### Knockdown of *ISWI1* affects cell survival and DNA elimination

We induced knockdown (KD) of *ISWI1* by feeding *Paramecium* with *ISWI1*- specific sequence triggering the cell’s internal RNAi machinery (Figure S1A). In a survival test of the post-autogamous (self-reproduction) progeny after *ISWI1*-KD over a period of three days, 86% of the cells did not survive beyond the first day after cells were fed again to resume vegetative division (Figure 1D). The remaining 14% of cells did not go through the usual rate of four vegetative divisions per day. In the control culture of *ND7*-KD (a gene required for exocytotic membrane fusion trichocyst discharge)(Skouri and Cohen 1997), the division rate of all the progeny remained unchanged, whereas in the positive control of *PGM-KD*, 90% of the cells did not survive as expected. In contrast to *ISWI1-KD*, for *ISWI5*-KD, 90% of the cells showed no substantial difference in division rate compared to the control cells (Figure S1B & S1C).

To test if the knockdown of *ISWI1* and *ISWI5* affect DNA elimination, we determined the retention status of germline-specific DNA elements in the newly developed MAC genome. First, we used probes against two well-known abundant families of transposons in the *Paramecium* MIC, Sardine and Thon. In *ISWI1-KD*, we could see greater retention of the Sardine and Thon, respectively compared to the control *ND7*- KD (Figure 1E). We further tested for IESs retention from a well-characterized locus using PCR with IES-flanking primers (Table T1). For the *ISWI1*-KD, all of the maternally-controlled (scnRNA-dependent) IESs tested by PCR were retained, as well as several of the non-maternally-controlled (scnRNA-independent) IESs (Figure 1F). For the *ISWI5*-KD, no retention of any of the IESs was observed (Figure S1D).

To focus our investigations on genome reorganization, all further experiments were, therefore, carried out for *ISWI1* only.

### ISWI1 is required for the complete excision of most IESs

To gain a genome-wide perspective on IES retention we performed high-throughput sequencing analysis of the genomic DNA isolated from the developing macronucleus from *ISWI1*-KD cell cultures (two biological replicates). As a control, we used genomic DNA from the developing macronucleus after *ND7*-KD (also a pair of biological replicates). IES retention scores (IRS) vary from 0.0 (complete IES excision) to 1.0 (complete failure of IES excision) upon the knockdown.

Approximately 35,000 (78%) IESs are sensitive to *ISWI1*-KD with a right-skewed retention score distribution (Figure 2A). IES retention scores of the biological replicates correlated well with each other (Pearson correlation coefficient: r=0.91). Generally, *ISWI1*-KD IES retention scores are modestly correlated with other known factors of excision machinery, correlating best with *DCL2/3/5*-KD (r=0.74) and *NOWA1/2*-KD (r=0.72; Figure S2A). *ISWI1*-KD retention score do not correlate as well with chromatin-related factors, *PTCAF1* (r=0.59) and *EZL1* (r=0.52).

**Figure 2.**
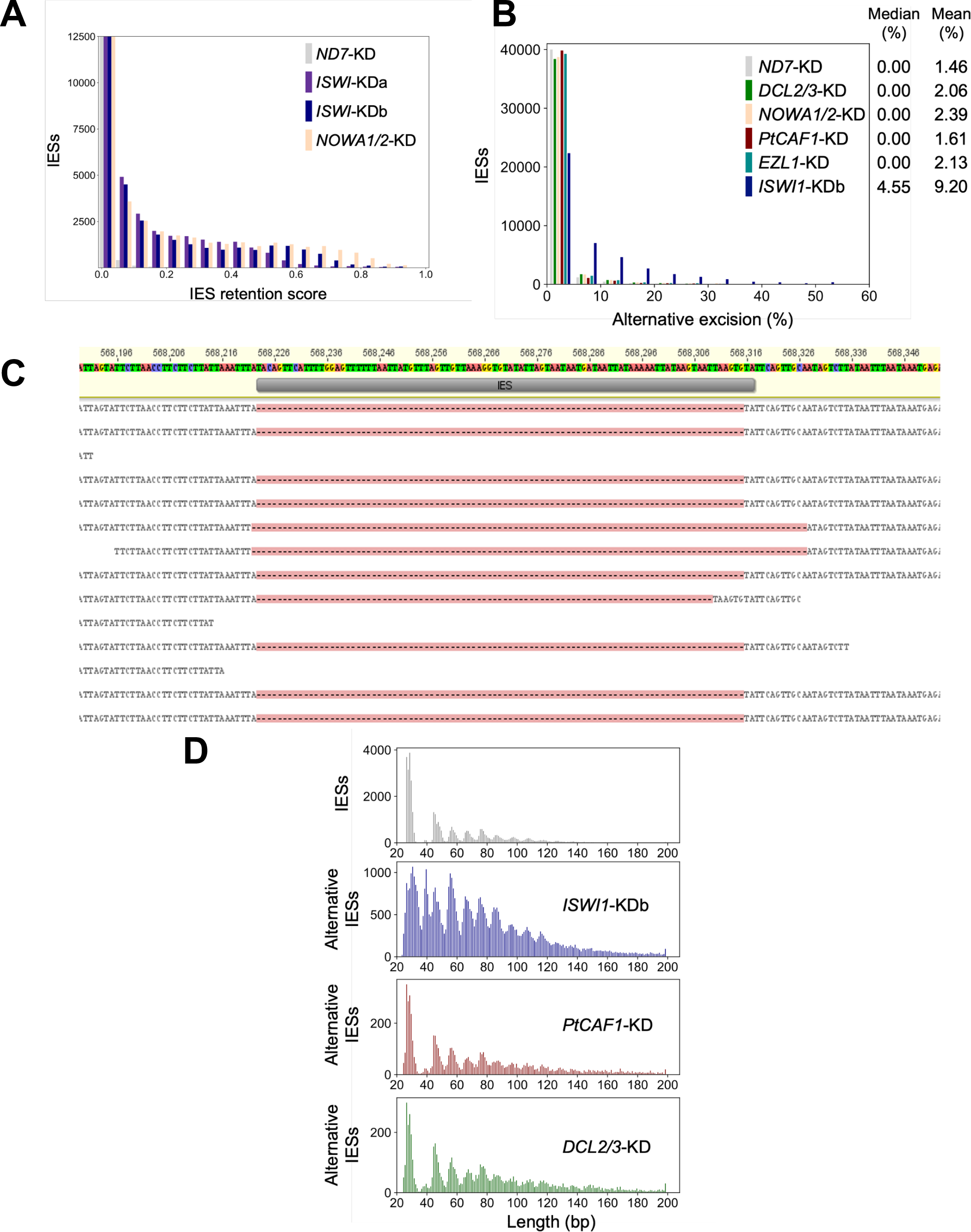
Genome-wide analysis of IES excision upon *ISWI1*-KD. (A) IES Retention Score (IRS) distributions for *ISWI1*-KD replicates. *ND7*-KD was used as a negative control. (B) Genome-wide analysis of alternative boundary excision in *ND7*-KD, *DCL2/3*-KD, *Nowa1/2*-KD, *Ezl1*-KD, *PTCAF1*- KD and *ISWI1*-KDb. Alternative excision (%) = 100* (alternative excised reads)/ (alternatively + correctly excised reads). (C) Reads mapped to an IES (IESPGM.PTET51.1.7.550914) showing both external (2 reads) and internal (1 read) alternatively excision; gaps opened in reads with excised IESs are indicated by dashes on a pink background. (D) Length distribution of conventional IESs compared to alternatively excised IESs in knockdowns of *ISWI1*, *PtCAF1* and *DCL2/3*.

As for most genes that influence IES excision, *ISWI1*-KD IES retention is length dependent. No periodicity of IES retention scores with respect to IES length is present (Figure S2B). Similar to other gene knockdowns, IES sub-terminal base frequency changes relative to IES retention scores for *ISWI1*-KD, i.e. base frequencies are relatively constant for the shortest and most common IESs but differ considerably in relation to IES retention score for longer IESs (Figure S2C (Swart et al. 2014)).

### *ISWI1-KD* enhances excision of IESs at alternative boundaries

Since ISWI homologs are involved in nucleosome positioning in other organisms, we sought to determine if *ISWI1*-KD would impact the precision in IES excision. Natural excision of IESs using alternative boundaries occurs at low frequency, impacting ∼16% of IESs in our negative control, *ND7-*KD (Figure 2B). In contrast, in *ISWI1-*KD, alternative boundary excision occurs at ∼65% of IESs (supported by one or more mapped reads; Figure 2C). This is substantially greater than for knockdowns of other genes necessary for IES excision, where the use of alternative IES boundaries is essentially the same as the control (Figure 2B). In general, though the amount of alternative IES excision for any given IES in *ISWI1-*KD is low (median 4.6%, mean 9.2%), it is substantially higher than that of other knockdowns (median 0%; mean 1.5-2.4%; Figure 2B).

In general, the length distribution of alternatively excised IESs, irrespective of the knockdown, follows a similar periodic pattern to normal IESs (Figure 2D), with smaller IESs more likely to result than larger ones (Figure 2D). Compared to normal IES excision, there is not as strong a preference for excision of the shortest IESs in alternative excision after *ISWI1-*KD. Interestingly, there are substantially more alternatively excised IESs in *ISWI1-*KD in the second, “forbidden” length peak around 35 bp than conventional IESs (Figure 2D). For conventional IESs, the suppression of this length peak, relative to its neighbours, is thought to reflect the inaccessibility of IESs (at this length) due to the conformational constraints of DNA and PGM dimerization (Arnaiz et al. 2012). We see a peak at this forbidden length in alternative excision events, regardless if they occur internally versus externally (Figure S3A and S3B). We do not observe a substantial increase of alternative IESs in the forbidden length range in other knockdowns. Thus, this appears to be a distinctive feature of *ISWI1*-KD. Furthermore, we propose that nucleosome positioning on or in proximity to IESs of this length plays a role in their excisability.

Cryptic IESs are IES-like sequences, which are randomly excised at low levels from DNA that is typically destined to become macronuclear during development(Swart et al. 2014). Since ISWI proteins play a role in repositioning nucleosomes in other organisms, we examined the effect of *ISWI1*-KD on cryptic excision. Cryptic excision in the *ISWI1-*KD was comparable to other knockdowns (Figure S3C and S3D). In other words, even if *ISWI1*-KD does alter nucleosomal positions, the net effect is not elevated cryptic IES excision.

### PTIWI01 and ISWI1 proteins interact *in vivo*

A C-terminal GFP fusion construct was made with *ISWI1* under the control of the putative *ISWI1* regulatory region, and linearized and injected into *Paramecium* vegetative macronucleus. When the early developing MACs were seen, using DAPI staining; the GFP signal of the fusion protein also accumulated in the developing MAC and remained there throughout the late developmental stages (Figure 3A).

**Figure 3.**
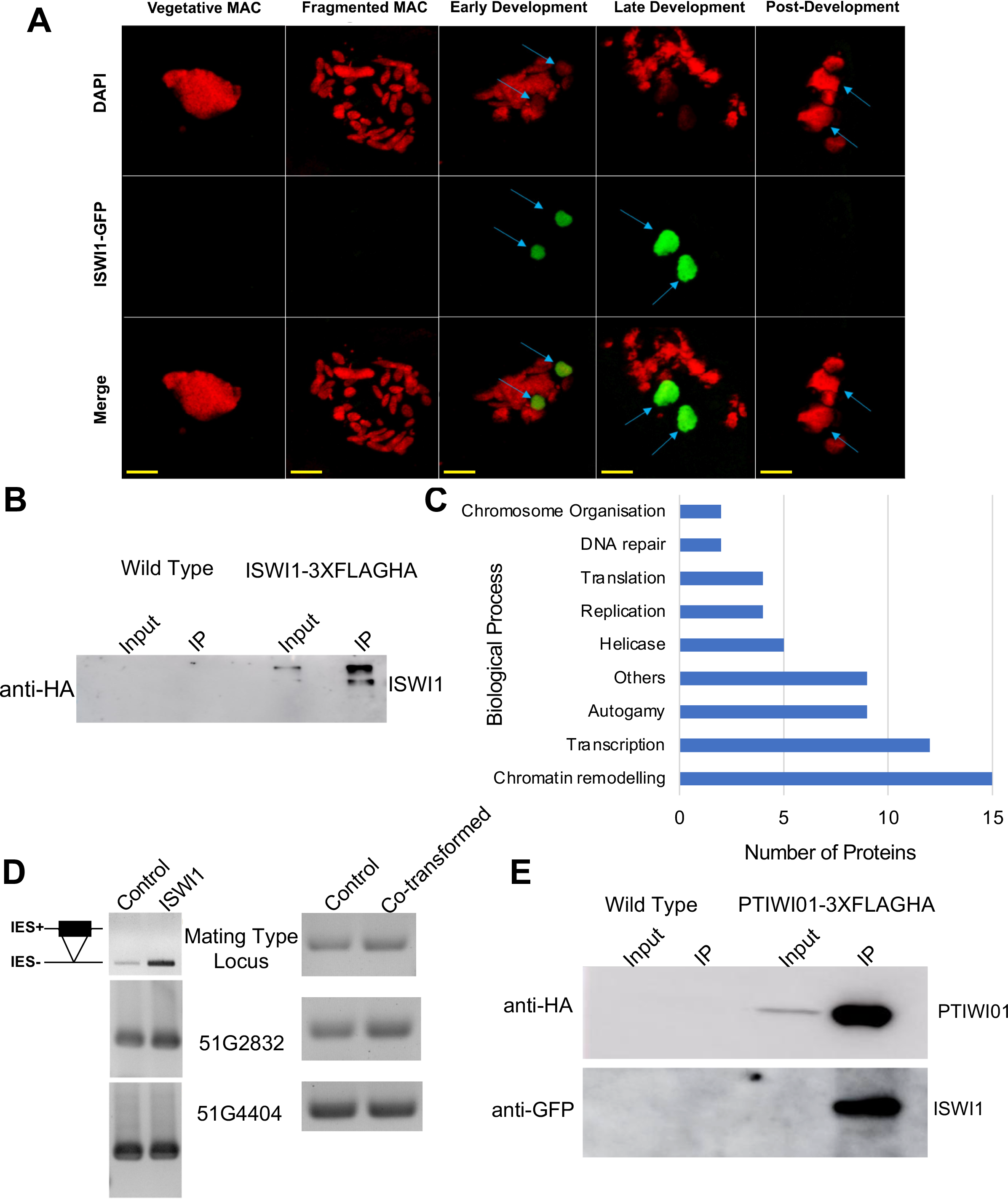
Localization, Co-immunoprecipitation and mass spectrometry analysis. (A) ISWI1-tagged C-terminally with GFP localizes in the developing MAC as soon as developing new MACs (panel Early Development ) become visible and remain there throughout late MAC development (panel Late Development). Red: DAPI, Green: *ISWI1*-GFP. Blue arrows identify developing MAC; scale bar 10µm. (B) Western blot analysis using anti-HA antibody after coimmunoprecipitation of ISWI-3XFlagHA fusion protein. Non-transformed cells (WT) of the same strain were used as the negative control. (C) Distribution of putative interacting partners into various categories according to their putative role in biological processes based on Paramecium DB search. (D) IES retention PCR (cropped inverted images). (E) Western blot analysis using anti-HA and anti-GFP antibodies after coimmunoprecipitation of PTIWI01-3XFlagHA fusion protein co-transformed with ISWI1-GFP. Non-transformed cells (WT) of the same strain were used as the negative control.

We sought to determine interacting partners of *Paramecium* ISWI1. First, we transformed *P. tetraurelia* cells with ISWI1 under its endogenous promoter and tagged with a 3XFlagHA at its C- terminal. We then co-immunoprecipitated (IP) ISWI1 to analyze the associated proteins by mass spectrometry. As a control, we performed the same experiment on non-transformed cells. Both controls and cells with the fusion protein were collected in two biological replicates during the developmental stage when ISWI1 localizes to the developing new MAC.

We effectively co-immunoprecipitated (IP) the fusion protein from the cell lysate. We detected a signal on Western blot using anti-HA antibody at the expected size of approximately (approx.) 124KDa (Figure 3B). The total IP samples were then analyzed by mass spectrometry (MS) where we could identify about 1500 proteins in total (Supplementary data D1). Among the 140 ISWI1-IP-exclusive proteins, we identified 9 proteins that are involved in *Paramecium* genome reorganization, notably PTIWI01, PTIWI09, NOWA1, and NOWA2 (Figure 3C, Supplementary Table T2).

Among the 9 characterized autogamy-specific proteins involved in genome reorganization, PTIWI01 was the most abundant among the ISWI1-IP replicates. Therefore, to check whether ISWI1 and PTIWI01 interact, we transformed *Paramecium* cells with 3XFLAGHA-tagged PTIWI01 and GFP-tagged ISWI1. We observed no growth defects or IES retention in the transformed cells either in single or co-transformed cells (Figure 3D). We succeeded in co-immunoprecipitating PTIWI01 (expected size approx. 90 KDa) at the developmental stage when ISWI1 is expressed (Figure 3E, upper panel). We then probed our IP samples with an antibody against GFP and detected a signal for ISWI1-GFP (expected size approx. 150 KDa), (Figure 4E, lower panel). Taken together, our data verify the interaction between ISWI1 and PTIWI01 in *Paramecium* (Figure 3E).

**Figure 4.**
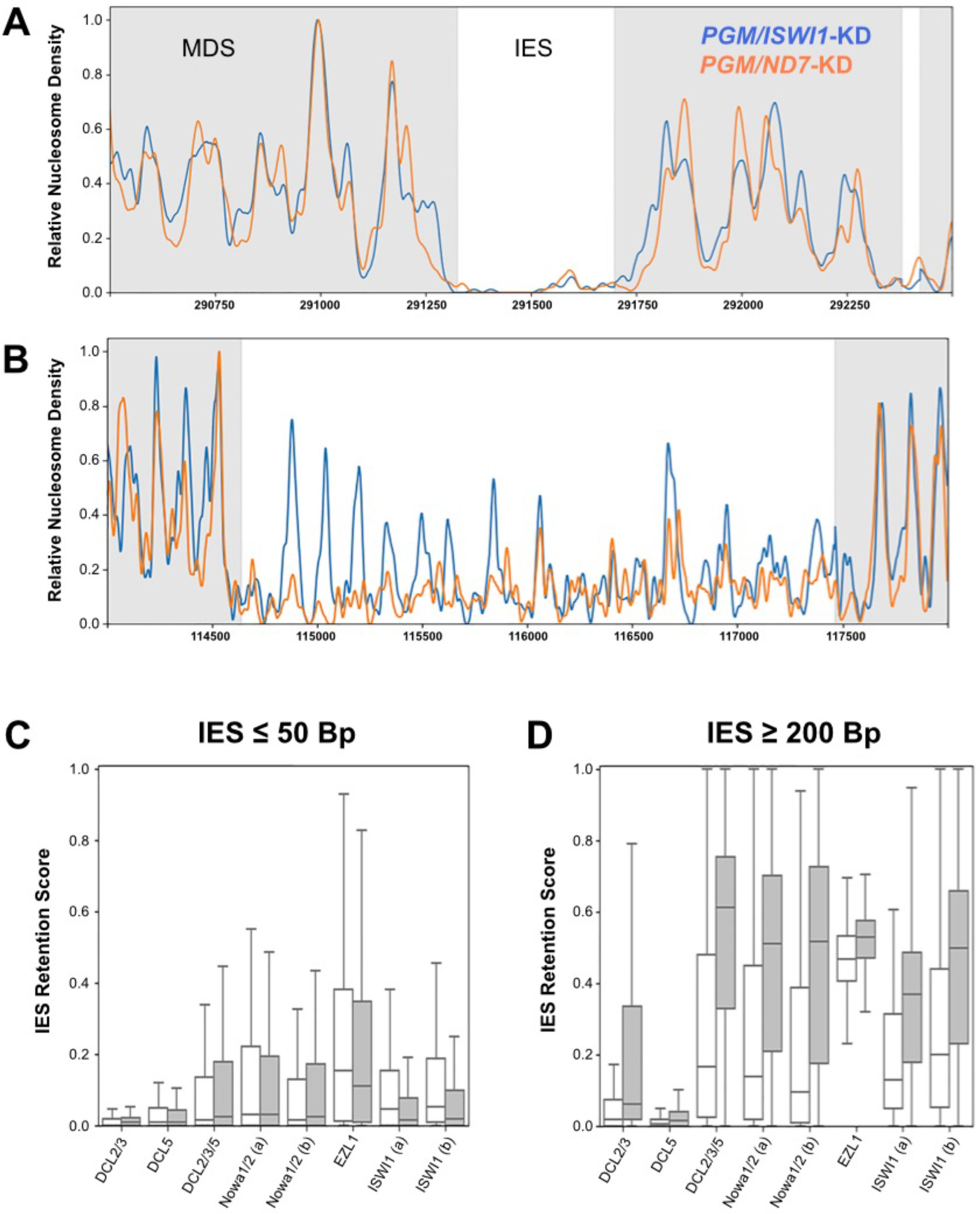
ISWI1-KD sensitive IESs are nucleosome rich. (A&B) Exemplar patterns of nucleosome density when *ISWI1* is present (orange) and absent (blue) in both *ISWI1* insensitive IESs (A; IESPGM.PTET51.1.24.274392; PGM/ND7 IRS=0.242, PGM/ISWI1-KD IRS=0.210) and *ISWI1* sensitive IESs (B; IESPGM.PTET51.1.144.109067; PGM/ND7 IRS= 0.251, PGM/ISWI1 IRS = 1.0). Note that nucleosome densities in IESs (white) are normalized to account for differences in IES retention, and that densities across MDS regions (grey) are for DNA from both the maternal and zygotic MAC. (C) Relationship between IES retention score distributions for the 10% most nucleosome poor in *PGM/ISWI1*-KD (white) and nucleosome rich (grey) IESs in different protein knockdowns for small (≤ 50 bp) IESs. (D) Relationship between IES retention score distributions for the 10% most nucleosome poor in *PGM/ISWI1*-KD (white) and nucleosome rich (grey) IESs in different protein knockdowns for large (≥ 200bp) IESs.

### *ISWI1*-sensitive IESs are nucleosome rich

We sought to determine whether nucleosome density changes occur around an IES during DNA elimination and whether this is ISWI1 dependent, we isolated developing macronuclear DNA from *PGM*-KD and *ISWI1/PGM*-KD cultures either with or without Atlantis dsDNAse treatment. Atlantis dsDNAse cleaves phosphodiester bonds in DNA and yields homogeneous populations of core nucleosomes. As *PGM* is a key component of the core endonuclease that cleaves IESs (Baudry et al. 2009; Bischerour et al. 2018; Arnaiz et al. 2012), which is also the last step of DNA elimination, we used *PGM*-KD as the control for our experiment, mapping the nucleosome density around IESs. A double knockdown of *ISWI1* with *PGM* is necessary to retain the majority of IESs to map the nucleosome density across them.

The distribution of retention scores in our *PGM/ND7*-KD are shifted skewed to the left (lower IES retention) compared to the reference *PGM*-KD silencing data sets (Swart et al. 2014; Arnaiz et al. 2012), whereas the IRS of *ISWI1/PGM*-KD is more similar to the knockdown expected for *PGM*-KD (Figure S4A). Previous experiments have shown that weakened IES retention due to dilution of gene knockdown can occur in *Paramecium* due to gene co-silencing (Arnaiz et al. 2012). Thus, the weaker silencing effect can be explained by the dilution of PGM silencing medium with ND7 silencing medium. This was done to ensure that the RNAi effects from both the *PGM/ND7* and the *ISWI1/PGM* knockdowns would be directly comparable.

Nevertheless, there are ∼5,500 IES shared between the *PGM/ND7-KD* and *ISWI1/PGM-KD* with nearly identical IRS (less than 0.05 difference) and another intermediate set of ∼6,000 IES with a difference in IRS, between the two co- silencings, ranging from ∼0.2 to ∼0.5. We attribute these differences largely to the knockdown of *ISWI1*.

To explore the impact of nucleosome density and *ISWI1* on IES retention, we defined large (≥ 200bp) IESs as being *ISWI1*-sensitive or *ISWI1*-insensitive. *ISWI1*- sensitive IESs are defined as those whose retention scores increase by ≥ 0.2 during *ISWI1/PGM* knockdown compared to only *PGM* knockdown, whereas *ISWI1*- insensitive IESs constitute the remainder. Afterwards, to account we normalized the nucleosome density of these IESs based on their IES retention scores, to account for differences in the efficacy of their excision. From these pools of IESs, we observe that *ISWI1*-insensitive IESs are typically depleted of nucleosomes whether *ISWI1* expression is suppressed or not (e.g., Figure 4A). However, IESs whose excision are sensitive to the presence of *ISWI1* clearly demonstrate local differences in nucleosome density compared to when *ISWI1* is depleted (Figure 4B).

Additionally, we observe a clear difference in the relationship between nucleosome density and IES retention associated with size (Figure 4C and 4D). Most IESs that are well below the size of a nucleosome (∼146 bp) are *ISWI1*-insensitive, with no clear difference in the impact of nucleosome density and their retention under numerous knockdowns (Figure 4C). However, the excision of longer IESs (≥ 200bp) with high nucleosome densities (top 10%) is strongly affected by the knockdown of key proteins involved in sRNA-mediated genome reorganization compared to those with low nucleosome densities (bottom 10%; Figure 4D).

### Impairment of the scnRNA pathway resembles *ISWI1-*KD

As NOWA1 and PTIWI01 were found in our ISWI1 immunoprecipitations (Supplementary Table T2), we sought to further determine how the early scnRNA pathway influences chromatin remodeling in *P. tetraurelia.* Since *NOWA1* is thought to mediate long ncRNA and scnRNA interaction necessary for IES excision (Nowacki et al. 2005; Sandoval et al. 2014; Swart et al. 2017), we chose to co-silence *NOWA1* and *PGM* for nucleosome profiling. Moreover, this strategy also helps us to avoid dilution of gene knockdown due to gene co-silencing of multiple genes which would have been the case if *PTIWI01*/*PTIWI09*/*PGM* knockdown was used.

As the nucleosome density surrounding most small IESs (≤ 100bp) was unclear in our *ISWI1-*-KD, arguably due to their sub-nucleosomal size, we chose to further focus on large (≥ 200bp) IESs and their nucleosome occupancy in *PGM-* KD/*NOWA1*. Comparisons of the most nucleosome-rich and -poor IESs in our *NOWA1/PGM*-KD show strikingly similar patterns to *PGM-*KD/*ISW1* for these large IESs. The most nucleosome rich IESs were sensitive to *PGM-*KD/*Nowa1* (*i.e.,* possess greater IES retention scores) compared to the most nucleosome-poor IESs (Figure S4B & S4C). Incidentally, these IESs show similar distributions of retention scores with *ISWI1*-sensitive IESs and other knockdowns demonstrably involved in the scan RNA pathway (Figure S4B & S4C).

## Discussion

*Paramecium* depends on efficient and accurate whole genome reorganization to produce a functional somatic nucleus during sexual reproduction. The excision of a third of IESs are sensitive to the presence of their counterpart sequences in the maternal macronucleus and require scnRNAs for their excision. However, the identification of numerous proteins required for the excision of both maternally-controlled and non-maternally controlled IESs (Arambasic et al. 2014; Lhuillier-Akakpo et al. 2014; Wasmuth and Lima 2017) suggests additional or alternative mechanisms beyond those envisaged in earlier models of RNA scanning and heterochromatin formation contributing to IES targeting and excision.

In this study, we have identified a putative ISWI, an ATP-dependent chromatin remodeler, that is required for the precise elimination of both maternally and non- maternally controlled IESs. ISWI proteins are highly conserved ATP-dependent chromatin remodelers (Corona et al. 1999) which regulate several biological processes (Yadon and Tsukiyama 2011), and now, as we have shown, also in genome editing. *ISWI1* is present in the developing macronucleus (Figure 3A) when the molecules responsible for genome reorganization cooperate to eliminate DNA.

Histone modification and heterochromatin formation is proposed to be a prerequisite for programmed DNA elimination in ciliates. The most evidence in support of this has been obtained for *Tetrahymena thermophila* (Liu et al. 2007; Xu et al. 2021). A similar model was proposed for IES excision in *Paramecium* as well (Coyne et al. 2012). It has been shown that histone modifications are required for targeting the excision of at least a subset of IESs ( (Ignarski et al. 2014; Lhuillier-Akakpo et al. 2014). Indeed, the knockdown of EZL1, a histone methyltransferase (Frapporti et al. 2019), affects the excision of the majority of IESs. Additionally, *EZL1*-KD affects the excision of IESs that are smaller than the size of a nucleosome (70.91% compared to 32.75% for *ISWI1*-KD, (Lhuillier-Akakpo et al. 2014)). Since heterochromatin regions generally spread across several kilobases in the genomes of other organisms (Margueron and Reinberg 2011; Huang et al. 2012), it was suggested that in *Paramecium*, H3K27me3 marks are placed locally (Lhuillier-Akakpo et al. 2014) .

Although it was recently shown that the transposable elements are enriched with these modifications (Frapporti et al. 2019), currently, there is no published information on H3K27me3 or H3K9me3 association with IESs. Moreover, H3K27me3 modification is not limited to the developing macronucleus and is also present in the fragments of the parental macronucleus (Frapporti et al. 2019; Ignarski et al. 2014; Lhuillier-Akakpo et al. 2014). This raises the possibility that inhibition of IES excision and the resultant cell lethality due to *EZL1*-KD and/or *PTCAF1*-KD may arise due to alteration in gene expression from the fragments during development.

Additionally, supporting the notion of indirect effects due to *EZL1*-KD, the nucleosome density of *EZL1*-sensitive and -insensitive IESs remains quite similar (Figure 4D). For *Paramecium* this is contrary to the conventional model of the requirement of heterochromatin for IES recognition and excision (Coyne et al. 2012). Thus, further experiments will be necessary to disentangle possible indirect effects from direct ones.

A subset of both maternally-controlled and non-maternally-controlled IESs were retained after the knockdown suggesting either a role for *ISWI1* in the joint machinery required for the excision of both classes of IESs or that *ISWI1* functions are agnostic to the different classes of IESs. Supporting the former, IES retention upon *ISWI1*-KD correlates modestly with *DCL2/3/5*-KD (as together they produce the sRNAs necessary to excise maternally controlled IESs; (Lepère et al. 2009; Sandoval et al. 2014); and large IESs (≥ 200bp) most sensitive to these knockdowns are substantially more nucleosome rich (Figure 4D). Upon *NOWA1-*KD we observe a similar impact on local nucleosome density for these large maternally-controlled IESs. Additionally, we also observe an interaction between PTIW01 and ISWI1 *in vivo* in our co-immunoprecipitation assay (Figure 3E). These data suggest that the sRNAs produced during genome remodeling in *Paramecium* confer some information regarding nucleosome spacing to *ISWI1*.

Uniquely among *Paramecium* proteins involved in IES excision investigated thus far, *ISWI1* gene silencing leads to elevated alternative IES excision (Figure 2B) suggesting that the endonuclease complex is not always able to correctly target the boundaries of an IES in the absence of *ISWI1*. The commonly accepted mechanism underlying ISWI function is that it controls the length of linker DNA and the chromatin architecture by altering nucleosome spacing (Corona et al. 2007; Xiao et al. 2001; Bartholomew 2014). We propose that the presence of nucleosomes on, or partially overlapping, an IES may be crucial for its targeting and accessibility to the excision machinery. Global nucleosome density changes are known to occur across genomes during cell lineage commitment as an additional regulatory mechanism (Erdel et al. 2011; Li et al. 2012).

Our results show, for longer IESs, those that are nucleosome-rich are more sensitive to *ISWI1*-KD and *NOWA1*-KD than those that are nucleosome-poor (Figure 4D and Figure S4C). This is in contrast to comparable sensitivity for nucleosome-rich and poor IESs in EZL1-KD and PtCAF1-KD. A plausible explanation could be that local nucleosome density changes are required to govern accessibility and possibly activating the endonuclease for DNA elimination. A similar explanation has been proposed for V(D)J recombination, where nucleosome location and occupancy changes were observed to regulate DNA recombination (Pulivarthy et al. 2016). We propose that once scnRNAs bind to IES complementary sequences in non-coding RNA ncRNAs (ncRNAs) in the developing new MAC, PTIWI01 relays the information to a ISWI1 complex that leads to local nucleosome density changes around nucleosome-rich IESs. Since the IES retention scores of *ISWI1*-KD are more strongly correlated with *DCL2/3/5-*KD and *NOWA1/2*-KD than with *DCL2/3*-KD and *PTIWI01/09*-KD iesRNAs bound to PTIWI10/11 likely interact in a similar manner with ISWI1 and IESs. The IESs are freed of nucleosomes and thus provide access and subsequent precise excision (Figure 5).

**Figure 5.**
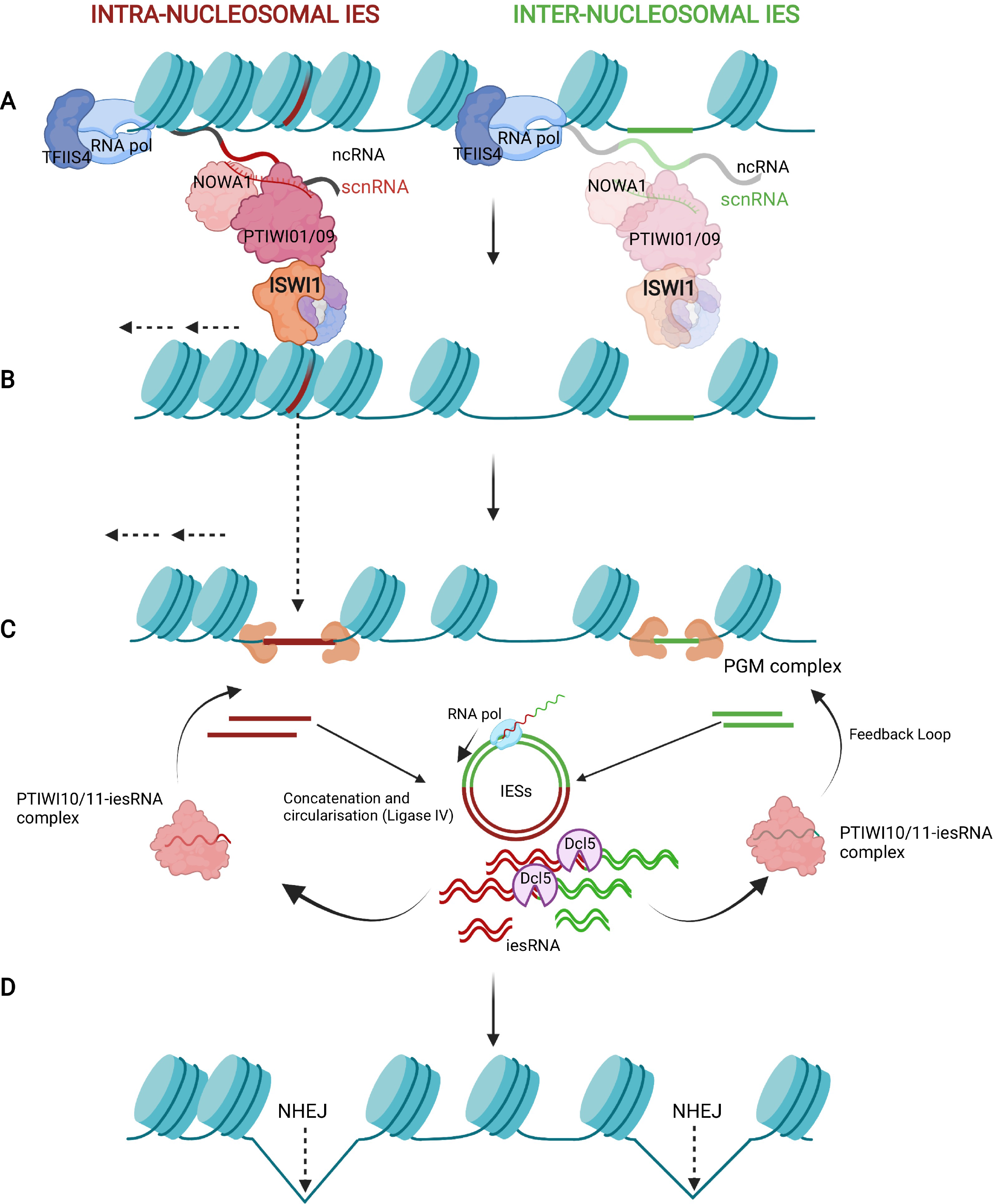
Proposed model of ISWI1 in elimination of germline-limited DNA. Scanning model during programmed DNA elimination in Paramecium tetraurelia created with BioRender.com. (A) Developing macronuclear genome is transcribed into non-coding RNA (ncRNA) by RNA polymerase assisted by TFIIS4. IES targeting scnRNA bound to PTIWI01/09 complex relays information to ISWI1 for the (B) repositioning of nucleosomes to make IES free of nucleosomes. (C) The repositioning of nucleosomes allow PGM complex to target the excision of the IESs. Excised IESs are concatenated by LIGASE IV into circular DNA that is then cleaved into iesRNA by dicer-like protein DCL5. iesRNA bound to PTIWI10/11 complex ensures the complete excision of IESs via a positive feedback loop. (D) After IES excision, chromosome ends are religated by non-homologous end joining (NHEJ) or is followed by chromosome fragmentation and de novo telomerization (not shown).

In conclusion, our findings reveal a role of *ISWI1* in targeting the germline-specific genomic sequences for their precise elimination, partially guided by scnRNAs which increase local chromatin accessibility through local chromatin remodelling, which ultimately leads to the formation of a functional somatic genome.

## Materials and Methods

### *Paramecium* Cultivation

Mating type 7 of Paramecium strain 51 were used in different experiments. Cells were cultured in Wheat Grass Powder (WGP; Pines International, Lawrence, KS) medium bacterized either with non-virulent *Klebsiella pneumoniae* or with *E. coli* and supplemented with 0.8 mg/l of β-sitosterol (567152, Calbiochem). Cells were either cultured at 27°C or at 18°C as per requirement. Clonal cell lines of *Paramecium* transformed with recombinant genes were maintained at 18°C as previously described (Beisson et al. 2010b).

### Silencing experiments, survival test and IES retention PCR

Silencing construct of *ISWI1* (Genbank accession: XM_001431568, XM_001431569) was made by cloning a 704bp construct from its C-terminal and cloned into an L4440 plasmid (using GGGTCTCACCTAAGATGAACG and TCACTTTCTTAACAGACTCAGATCC). For *ISWI5* (Genbank accession: XM_001432642), silencing construct was made by cloning a 1106bp long fragment into L4440 plasmid (using ATGAGTGAAAGTGAAGATGAG and AGATTTCGTCCTTCTTAACAT). The plasmids were then transformed into HT1115 (DE3) *E. coli* strain. Cells were seeded into the silencing medium at a density of 100 cells/ml and silencing was carried out according to previously described protocol (Beisson et al. 2010d). After the cells finished autogamy, 30 post-autogamous cells were transferred individually to three-well glass slides containing medium bacterized with avirulent *K. pneumoniae* for the survival test. Cells were monitored for three days (approximately 12 divisions) and categorized into three groups according to their observed phenotype. In parallel, a 100ml culture was harvested for DNA extraction using GeneElute –Mammalian Genomic DNA Miniprep Kit (Sigma- Aldrich). PCRs were done on different genomic regions flanking an IES.

### Dot Blot

Dot blot assays were conducted following standard protocols (Brown 2001). Briefly, 3 μg of DNA from post-autogamous cultures were blotted onto a nylon membrane (Hybond N+ XL membrane, Amersham). Probes specific to Sardine and Thon transposons and actin (first 240bp of the gene) labelled with α-32P dATP (3000 Ci/mmol) using RadPrime DNA Labeling System (Invitrogen) were used. The signal was quantified with ImageJ 1.48e.

### Northern Blot

10 μg of RNA were run in a 1.2% agarose denaturing gel and transferred to a nylon membrane (Hybond N+ XL membrane, Amersham) by capillary blotting. After transfer, the membrane was crosslinked twice with UV (120000µJ/cm^2^). Specific probes labelled with α-32P dATP (3000 Ci/mmol) using RadPrime DNA Labeling System (Invitrogen) for ISWI1, ISWI5 and rRNA were used for hybridization.

Membranes were screened using the Typhoon Imaging system (GE Healthcare).

### GFP tagging, microinjection and GFP localization experiment

A set of specific *ISWI1* specific primers (5’- GTA GAA TCC TAT TGA TAG GAG GAG-3’ and 5’-TGG CTC TAA GAA ATT CAT TTA T-3’) were used for the amplification full gene including 227 bp upstream and 62 bp downstream of the coding region. *ISWI1* was tagged with GFP on its C-terminus. The construct was linearized using NaeI restriction enzyme (R0190S, New England Biolabs) and injected into the macronucleus of the vegetative cells as previously described (Beisson et al. 2010a). Cells positive for GFP expression were collected during different stages of autogamy and either stored with 70% ethanol at -20°C or directly fixed with 2% PFA in PHEM and then washed in 5% BSA with 0.1% Triton X-100.

Cells were then counterstained with DAPI (4,6-diamidino-2-2phenylindole) in 5% BSA with 0.1% Triton X-100 and mounted with Prolong Gold Antifade mounting medium (Invitrogen). Images were then acquired with Olympus Fluoview FV1000 confocal microscope system with PLAPON 60X O SC NA 1.40. Images were analysed and given pseudo-colour on Imaris software.

### Immunoprecipitation and mass spectrometry

*ISWI1* specific primers (5’- GTA GAA TCC TAT TGA TAG GAG GAG-3’ and 5’-TGG CTC TAA GAA ATT CAT TTA T-3’) were used for the amplification of the full gene with regulatory regions. The gene was tagged with 3XFLAGHA at its C-terminal.

4.5X10^5^ cells were harvested and crosslinked with 1%Paraformaldehyde by incubating for 10 minutes (min) at room temperature. Cells were then quenched using 100µl of 1.25M Glycine and incubated at room temperature for 5 min. Cells were washed once with PBS for two minutes at 500Xg. Further steps were carried at on ice and/or at 4°C. 2ml of lysis buffer (50mM Tris pH 8.0, 150mM NaCl, 5mM MgCl2, 1%Triton X100, 1X Protease inhibitor (Roche,11836170001), 10% glycerol) was added and the cells were sonicated (Branson Digital Sonifier) with 55% amplitude for 15 seconds. The lysate was then centrifuged for 30 minutes at 13’000Xg or until the lysate was clear. 50µl of bead slurry (HA High Affinity Matrix,11815016001, clone 3F10, Roche) was used per IP sample and was washed thrice by centrifuging for 2min at 500Xg. After washing the beads, 1ml of the lysate was mixed to the beads and incubated overnight with agitation at 4°C. After the incubation, the beads were washed five times with the IP buffer (10mM Tris pH8.0, 150mM NaCl, 0.01% NP-40, 1mM MgCl2, 1X Protease inhibitor (Roche,11836170001), 5% Glycerol) for 2min at 500Xg. NP-40 was added freshly in the buffer. Proteins were then eluted by adding 50µl of the 2X loading buffer (10%SDS, 0.25M Tris ph6.8, 50% Glycerol, 0.2M DTT, 0.25% Bromophenol blue).

Mass spectrometry analysis was done at the Proteomics & Mass Spectrometry Core Facility (PMSCF), University of Bern. The mass spectrometry proteomics data have been deposited to the ProteomeXchange Consortium (Deutsch et al. 2020) via the PRIDE (Perez-Riverol et al. 2019) partner repository with the dataset identifier PXD027206.

For co-transformation with ISWI1-GFP, PTIWI01 with primers in its regulatory regions was (CATTTTTAAGAGATTTCAATAAAACAATTATCC and GTGCTTTGAAAATCAATGAAAATCA) amplified and 3XFLAGHA was fused at its N- terminal. After linearisation with NaeI, both constructs were mixed in equal proportion for microinjection. Co-immunoprecipitation assay was performed as explained above with a slight modification. Sonication was done with 52% amplitude for 20 seconds using MS72 tip on Bandelin Sonopulse.

### Western Blot

Western blot on IP samples was done by running a 10% SDS-PAGE gel, and the proteins were transferred on 0.45μm nitrocellulose membrane (10600002 Amersham, GE Healthcare). The membrane was blocked with 5% BSA in PBS for one hour at room temperature. The membrane was then incubated overnight at 4°C with anti-HA (sc805, Santa Cruz, RRID:AB_631618 ) in a dilution of 1:500. A goat anti-rabbit HRP conjugate (sc2004, Santa Cruz, RRID:AB_631746) in a dilution of 1:5000 was used after washing the membrane with PBS/0.1% Tween-20 for 10 minutes (three times). For PTIWI01-3XFLAGHA IP, the membrane was incubated with either anti-HA (sc-7392 HRP, Santa Cruz, RRID:AB_627809) in a dilution of 1:500 or with anti-GFP (ab290, Abcam, RRID:AB_303395) in a dilution of 1:1000.

The secondary antibody incubation was done for 1h at room temperature and the membrane was washed thrice with PBS/0.1% Tween-20 for 10 minutes. The membrane was then washed once for 5 minutes with 1X PBS before imaging. The membrane was scanned using chemiluminescence settings on an Amersham Imager 600 (GE Healthcare).

### Phylogenetic analyses

ISWI proteins were identified (OG5_127117) and retrieved using PhyloToL (Cerón-Romero et al. 2019). Briefly, multi-sequence alignments were constructed using MAFFT (Katoh and Standley 2013) and then iteratively refined with GUIDANCE2 (Sela et al. 2015), which identifies and removes spurious sequences and columns, preserving phylogenetically informative regions in the alignment. This refined alignment was then passed to RAxML (Stamatakis 2014), and used to generate 200 bootstrap replicates.

### Macronuclear isolation and Illumina DNA-sequencing

The samples for MAC isolation were collected from *ND7-KD, ISWI1-KD,* and *PTCAF1-KD* cultures from the cultures three days post autogamy as described previously (Arnaiz et al. 2012). Paired-end libraries (Illumina TruSeq DNA, PCR-free) were made according to the standard Illumina protocol. Library preparation and sequencing was done at the NGS platform, University of Bern.

### Reference genomes

The following reference genomes were used for analysing DNA-seq data. MAC: http://paramecium.cgm.cnrs-gif.fr/download/fasta/ptetraurelia_mac_51.fa MAC+IES: http://paramecium.cgm.cnrs-gif.fr/download/fasta/ptetraurelia_mac_51_with_ies.fa

### IES retention and alternative boundary analysis

IES retention scores were calculated with the MIRET component of ParTIES (Denby Wilkes et al. 2016). IES retention scores are provided as Supplementary Data D2 (ISWI1_MIRET.tab).

The MILORD component (default parameters) of a pre-release version (13 August 2015) of ParTIES was used to annotate alternative and cryptic IES excision. For each IES with alternative or cryptic excision, the identifiers for the supporting reads are recorded. Output for this is provided as Supplementary Data D3- (CAF1_MILORD.gff3.gz, DCL23_MILORD.gff3.gz, ISWI1-b_MILORD.gff3.gz, ND7-b_MILORD.gff3.gz, NOWA1_MILORD.gff3.gz). IRS correlations, relationship of IRS with length and sub-terminal frequencies were calculated as described previously (Swart et al. 2014). IES retention scores for PGM/ISWI1-KD and PGM/ND7-KD are provided in Supplementary Data D4 (PGM_ND7_ISWI_MIRET.tsv).

### Nucleosomal DNA Isolation and Illumina DNA-sequencing

Macronuclear DNA isolation protocol was followed up to the stage of ultracentrifugation. After ultracentrifugation, the pellet containing macronucleus was washed twice with chilled 1xPBS pH 7.4, and the excess PBS was removed by centrifuging at 200g for 2 minutes at 4°C. All the steps from here were optimized from the standard protocol provided with the EZ Nucleosomal DNA Prep Kit (D5220, Zymo Research). Briefly, 1mL of chilled Nuclei Prep Buffer was used to resuspend the cell pellet before incubating on ice for 5 minutes. The nuclear pellet was then centrifuged at 200g for 2 minutes at 4°C. After washing twice with Atlantis Digestion buffer, the pellet was resuspended in 1ml of Atlantis Digestion Buffer. 500µl of the reaction was then used for DNA isolation without digestion as a control. The remaining 500µl of the reaction was used for nucleosomal DNA isolation and 35µL of the Atlantis dsDNAse. The reaction was incubated at 42°C for 20minutes. After 20 minutes, the reaction was stopped by adding MN Stop Buffer and the nucleosomal DNA isolation was carried out according to the kit protocol (D5220, Zymo Research). Illumina TruSeq PCR free DNA library was prepared without bead-based size selection followed by a preparative size selection on the PippinHT to remove non- ligated adaptors and library molecules with inserts >500 bp.

### Nucleosome Density Measurements

Raw read pairs generated from the Atlantis dsDNAse digested samples (*PGM- KD/ND7*-KD, *PGM/ISWI1*-KD, and *PGM/NOWA1*-KD) were mapped against the *Paramecium tetraurelia* MIC genomes (Arnaiz et al. 2012; Guérin et al. 2017) using Bowtie2 (Langmead and Salzberg 2012) with standard parameters, permitting a single mismatch. From the outputs, those read pairs with an insert size corresponding to a single nucleosome (*i.e.,* 146 bp ± 20 bp) were kept and then calculated the per-bp coverage of the nucleosome. These data were normalized using the TPM to the genomic scaffolds to reduce impacts from stochasticity in sequencing depth. Following normalization, these data were then further smoothed with a Gaussian filter (standard deviation = 10 bp).

Germline genome regions to analyze were selected based on patterns between the IES retention scores of the *PGM/ND7* and the experimental, *PGM/ISWI1 and PGM/NOWA1*, knockdowns. All IESs analyzed had retention scores in the *PGM/ND7*-KD ≥ 0.2. This minimum threshold was chosen to increase the likelihood that the observed nucleosome occupancy is attributable to the developing somatic genome, rather than the corresponding germline loci and/or differences in accessibility due to IES excision (no signal would be observed in completely excised IESs). From this pool of IESs, we defined IESs as *ISWI1/NOWA1*-sensitive or *ISWI1/NOWA1*-insensitive. Sensitive IESs are those with an observed increase in retention score (≥ 0.2) in the experimental knockdowns compared to the control *PGM/ND7-*KD, whereas the rest were classified as insensitive IESs. We further normalized the nucleosome density of these IESs based on the ratio of their IES retention scores in *PGM/ND7*-KD and *PGM/ISWI1*-KD to further account for differences in IES excision efficiency. These normalized IESs and their surrounding genomic regions were then identified for further analyses.

### Data availability

All raw sequencing data are available at the European Nucleotide Archive under the accession number PRJEB21344. Accession numbers for individual experiments are as follows; ERR2010817 for *ISWI1*-KD(a), ERR2010816 for *ISWI1*-KD(b), ERR2010818 for *PTCAF1*-KD, ERR2010819 for *ND7*-KD, ERR2798685 for *PGM/ND7*-KD DNA, ERR2798686 for *PGM/ISWI1*-KD, ERR2798687 for *PGM/ISWI1*-KD nucleosomal DNA, ERR2798688 for *PGM/ND7*-KD nucleosomal DNA.

### Competing financial or non-financial interests

The authors declare no competing financial or non-financial interests.

## Acknowledgements

We are thankful to Dr. Nasikhat Stahlberger for her technical support and to all the members of the Nowacki lab for their suggestions and fruitful discussions.

This research was supported by grants from the European Research Council (ERC): 260358 “EPIGENOME” and 681178 “G-EDIT”, Swiss National Science Foundation: 31003A_146257 and 31003A_166407 and from the National Center of Competence in Research (NCCR) RNA and Disease to MN, and the Max Planck Society (ES).

## Author Contributions

MN conceived the project, provided guidance and edited the manuscript. AS performed the experiments. SG did macronuclear isolation of the *ISWI1-KDa* sample. TS performed NOWA1 experiments and cloned PTIWI01-3X FLAGHA. MI cloned the RNAi construct for ISWI1. AS, XXMA and ECS wrote the manuscript. AS, ECS and XXMA performed the bioinformatic analyses.

## Materials & Correspondence

Correspondence and material requests should be addressed to Mariusz Nowacki.

---

**Supplementary Figure S1.**
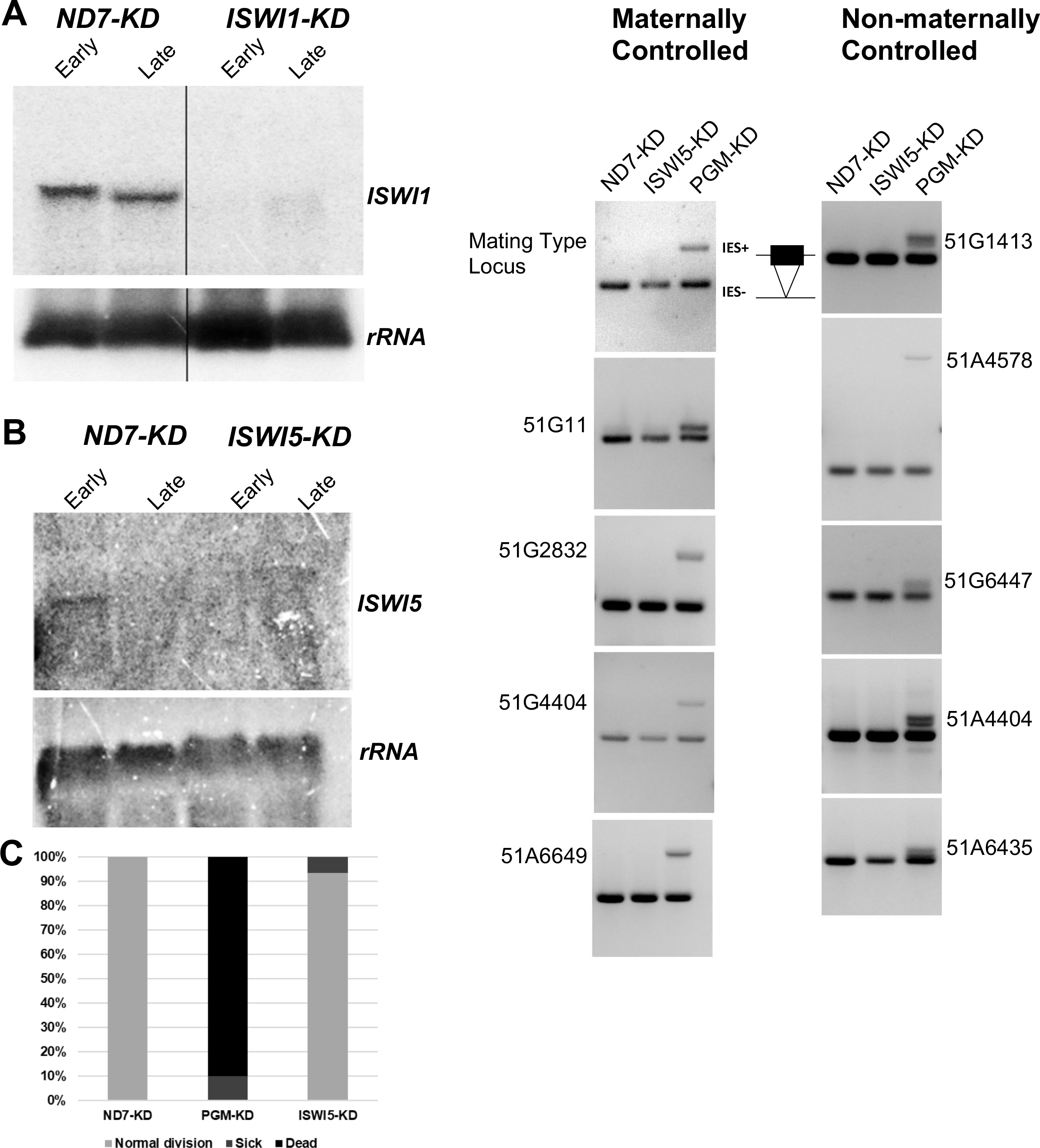
(A) & (B) Northern blot analysis using *ISWI1-*specific and *ISWI5*-specific probes respectively. rRNA probe was used as a loading control against ribosomal RNA. Early: ∼50% of cells with fragmented parental macronucleus; Late: the majority of cells with a visible anlagen. *ND7*-KD is used as a control to confirm mRNA expression. (C) Survival test graph. Dead cells are represented in black, sich in dark grey and cells diving at a normal rate in light grey. *PGM-KD* is used as a positive control and *ND7-KD* as a negative control. (D) IES retention PCR (cropped inverted images). Five maternally controlled IES and five non-maternally controlled IESs are shown. The IES+ band represents retained IES; the IES-band represents excised IES; additional bands are likely PCR artefacts or primer dimers.

**Supplementary Figure S2.**
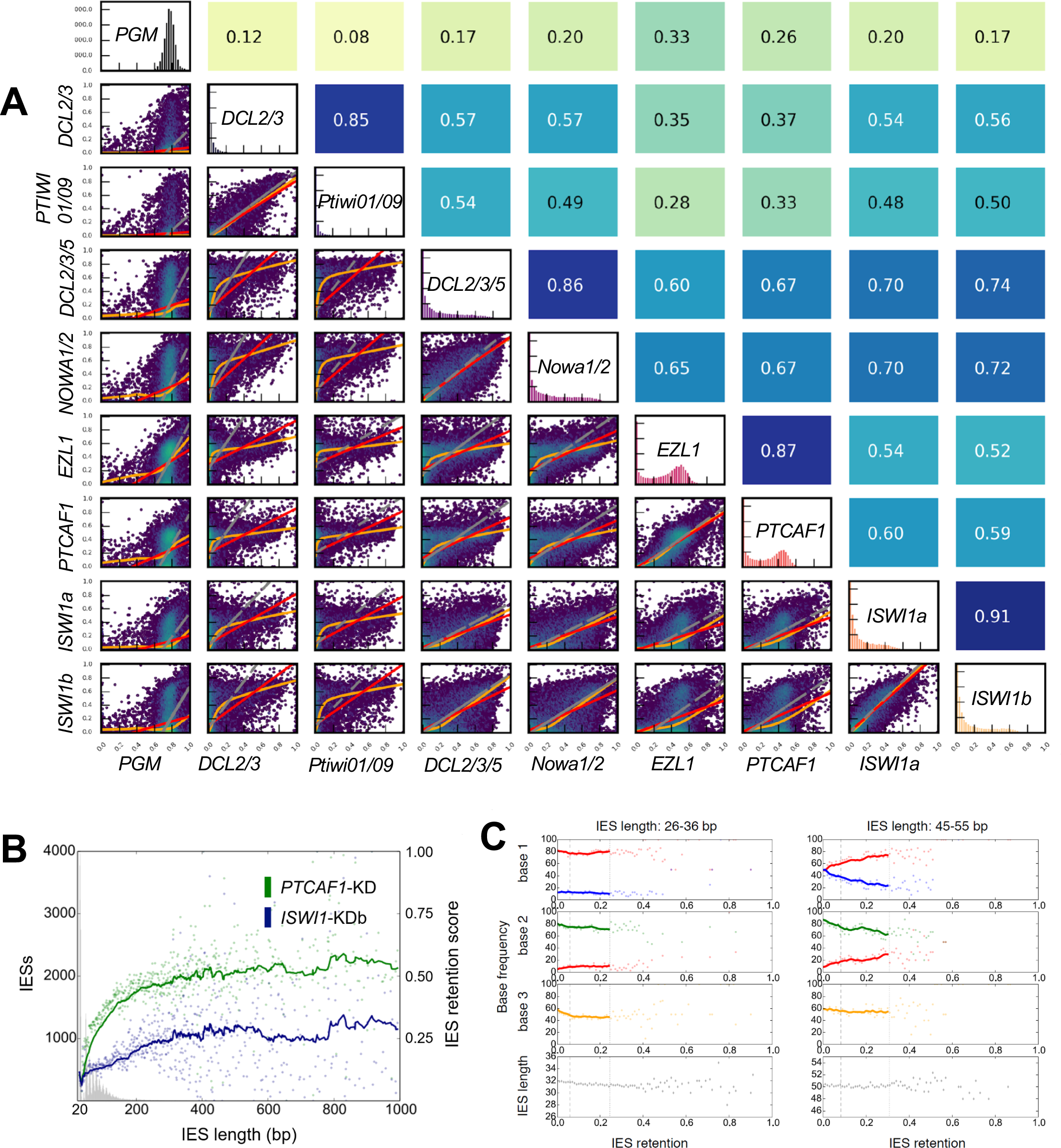
(A) Relationships in IES retention among knockdown pairs. Hexagonal binning of IES retention scores was used to generate the plots. Pearson’s correlation coefficients are given above each subgraph. Red lines are for ordinary least-squares (OLS) regression, orange lines for LOWESS, and grey lines for orthogonal distance regression (ODR). (B) IRSs versus IES length as described previously. (C) Base frequencies of the first three bases after the TA repeat relative to the IRS of ISWI1-KDb from the first and third Paramecium tetraurelia IES length peak.

**Supplementary Figure S3.**
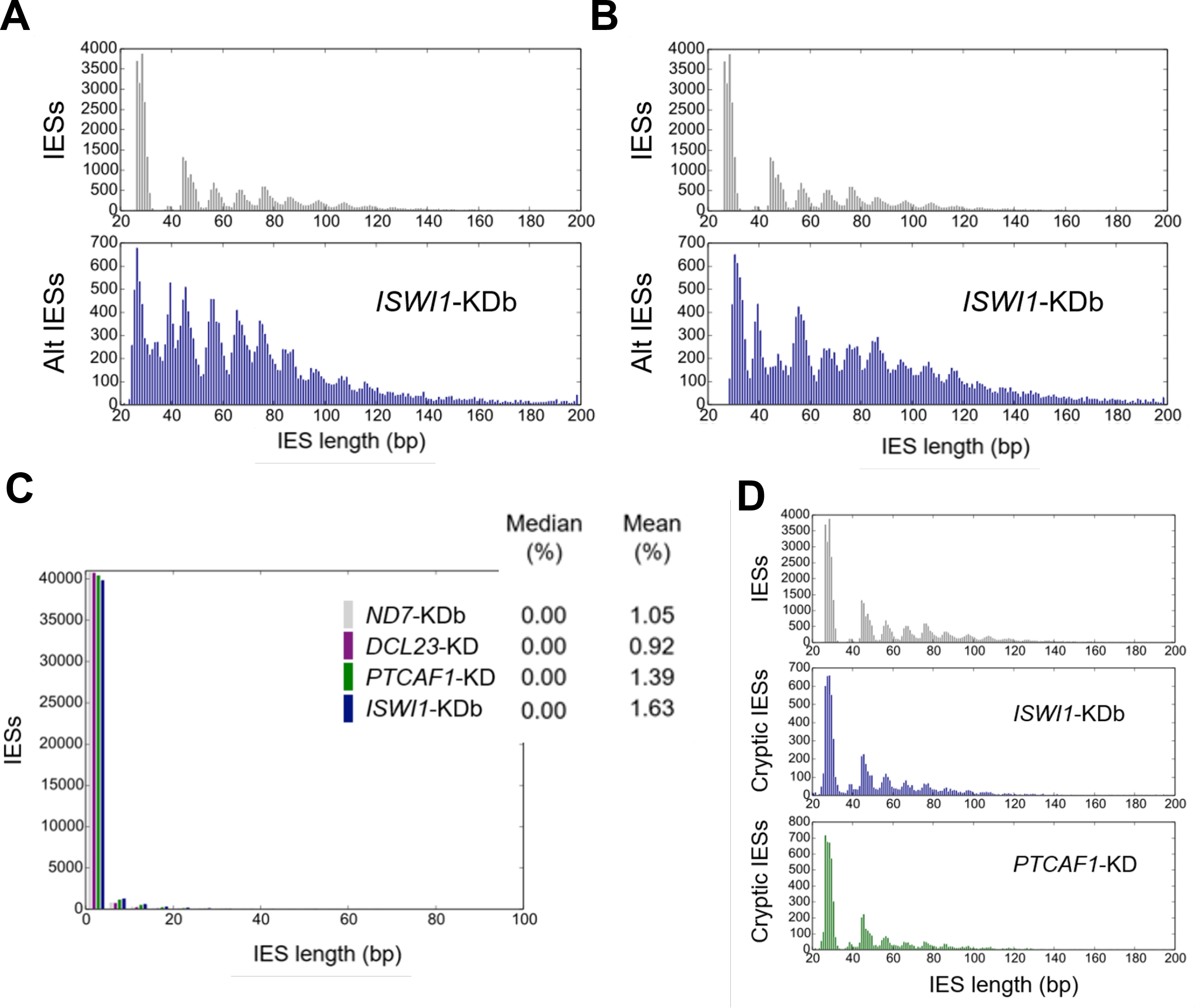
(A) Length distribution of internally excised alternative (Alt) IES boundaries (B) Length distribution of externally alternative (Alt) excised IES boundaries respectively. (C) Genome-wide analysis of cryptic IES excision. Cryptic excision (%) = 100 * (cryptically excised reads) / (all reads). (D) Length distribution of cryptically excised IES.

**Supplementary Figure S4.**
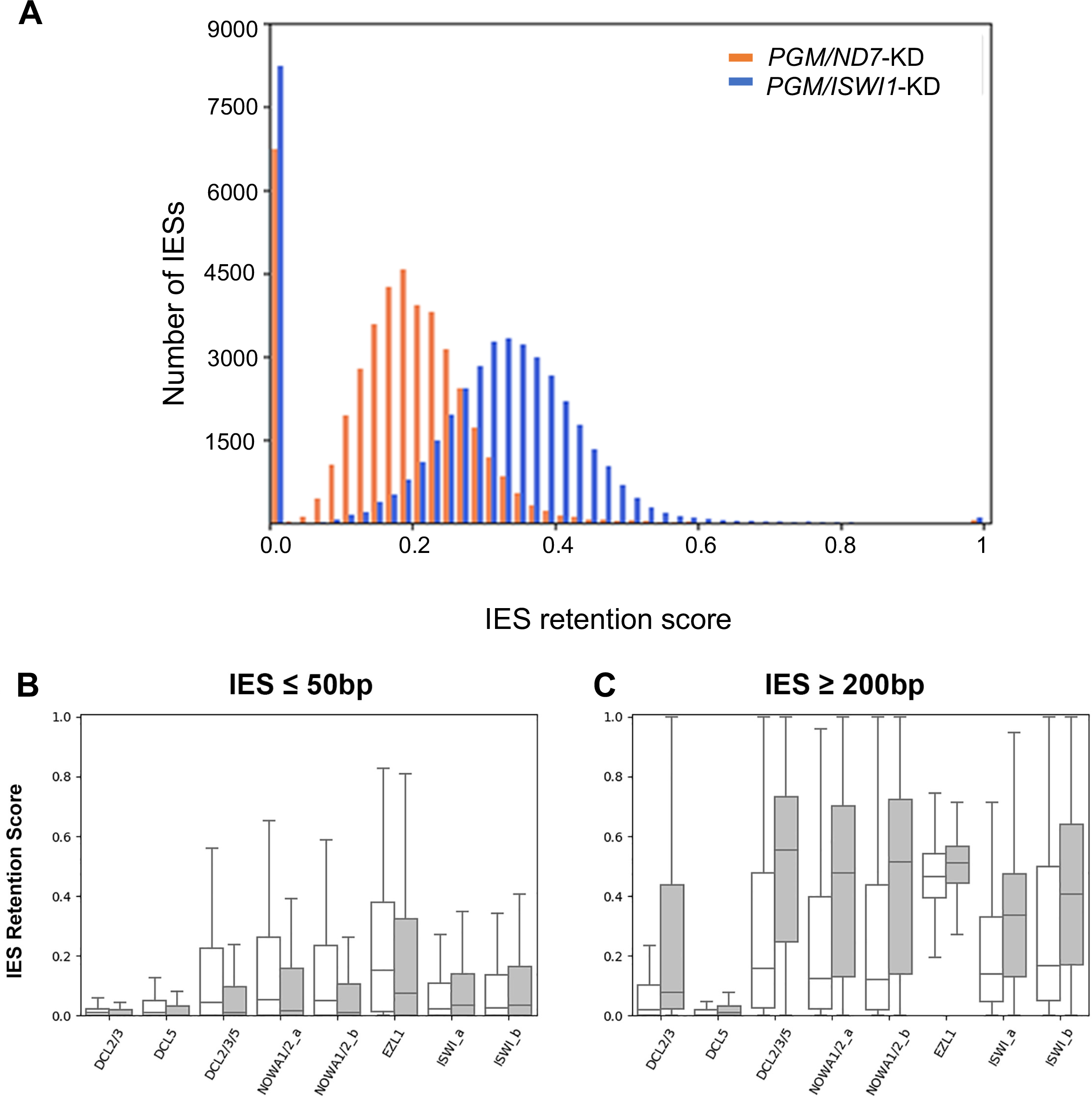
(A) IES Retention Score (IRS) distributions for *PGM/ND7*-KD and *PGM/ISWI1*-KDs. (B) & (C) Relationship between IES retention score distributions for the 10% most nucleosome poor in *PGM/NOWA1*-KD (white) and nucleosome rich (grey) IESs in different protein knockdowns for (A) small (≤ 50 bp) IESs and (B) large (≥ 200bp) IESs.

**Supplementary Table T1.**
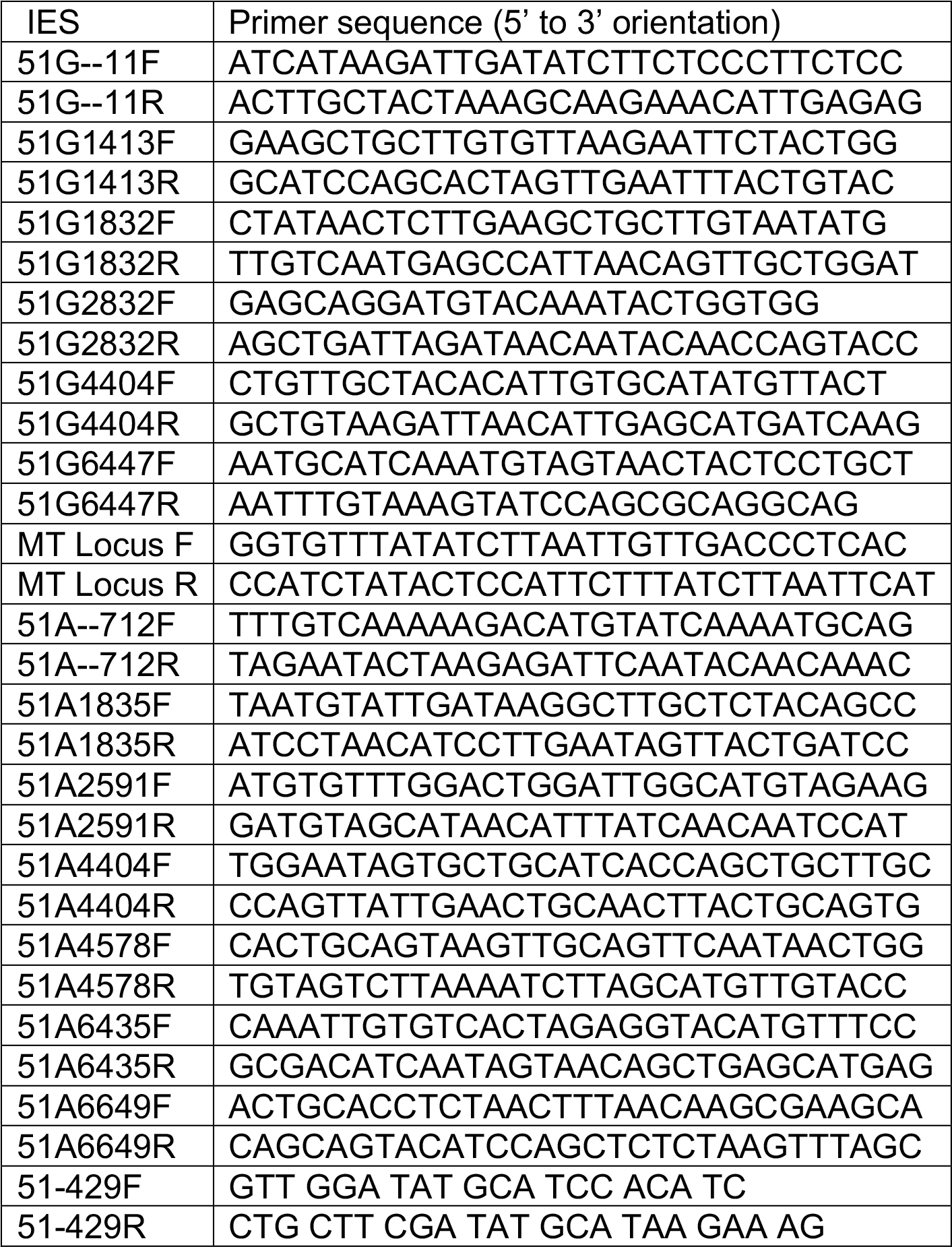
Primers used in IES retention PCRs. F: forward primer; R: reverse primer.

**Supplementary Table T2.** Mass spectrometry analysis of ISWI1-3XFLAGHA co-immunoprecipitation (IP). Majority protein IDs correspond to the *Paramecium* Database (Arnaiz and Sperling, 2011) accession numbers of the proteins identified by MS. MS/MS count ISWI1 & MS/MS count ISWI1 represents total peptide count in ISWI1- 3XFLAGHA IP replicates. MS/MS count WT1 & MS/MS WT2 represents total peptide count in negative control to IP. MS/MS count represents combined peptide counts in the replicates.

| Majority protein IDs | MS/MS<br>count<br>ISWI1 | MS/MS<br>count<br>ISWI2 | MS/MS<br>count<br>WT1 | MS/MS<br>count<br>WT2 | MS/MS<br>count | Entrez_Protein |
| --- | --- | --- | --- | --- | --- | --- |
| PTET.51.1.P0140243 | 243 | 261 | 0 | 0 | 504 | XP_001431605 |
| PTET.51.1.P0440186 | 90 | 104 | 0 | 0 | 194 | XP_001447805.1 |
| PTET.51.1.P0140243 | 58 | 68 | 0 | 0 | 126 | XP_001431606 |
| PTET.51.1.P1370127 | 62 | 61 | 0 | 0 | 123 | XP_001431060 |
| PTETP2700007001 | 61 | 50 | 0 | 0 | 111 | XP_001441411 |
| PTETP300037001 | 49 | 45 | 0 | 0 | 94 | XP_001442677 |
| PTET.51.1.P0420126 | 42 | 49 | 0 | 0 | 91 | XP_001447124 |
| PTETP7100004001 | 26 | 43 | 2 | 3 | 74 | XP_001456124 |
| PTET.51.1.P0180124 | 33 | 36 | 0 | 0 | 69 | XP_001437349 |
| PTET.51.1.P0070284 | 39 | 28 | 0 | 0 | 67 | XP_001455634 |
| PTET.51.1.P0480005 | 34 | 29 | 2 | 0 | 65 | XP_001448777 |
| PTETP10500011001 | 31 | 33 | 0 | 0 | 64 | XP_001424861 |
| PTET.51.1.P0110326 | 19 | 26 | 3 | 8 | 56 | XP_001425954 |
| PTET.51.1.P0330075 | 26 | 20 | 0 | 0 | 46 | XP_001443869 |
| PTET.51.1.P0560063 | 17 | 22 | 2 | 2 | 43 | XP_001451492 |
| PTET.51.1.P0720184 | 19 | 20 | 0 | 1 | 40 | XP_001456414 |
| PTET.51.1.P0990120 | 17 | 22 | 0 | 0 | 39 | XP_001462544 |
| PTETP13700005001 | 19 | 17 | 0 | 0 | 36 | XP_001431029 |
| PTET.51.1.P0040036 | 18 | 17 | 0 | 0 | 35 | XP_001446004 |
| PTET.51.1.P1720081 | 19 | 15 | 0 | 0 | 34 | XP_001436697 |
| PTET.51.1.P0610231 | 17 | 16 | 0 | 0 | 33 | XP_001453395 |
| PTET.51.1.P0380048 | 17 | 15 | 0 | 0 | 32 | XP_001445418 |
| PTET.51.1.P0200322 | 14 | 16 | 0 | 0 | 30 | XP_001439097 |
| PTET.51.1.P1060046 | 11 | 16 | 0 | 0 | 27 | XP_001425020 |
| PTET.51.1.P0030037 | 9 | 15 | 2 | 1 | 27 | XP_001442334 |
| PTET.51.1.P1600114 | 9 | 16 | 0 | 0 | 25 | XP_001435111 |
| PTET.51.1.P0470202 | 10 | 15 | 0 | 0 | 25 | XP_001448725 |
| PTET.51.1.P0410183 | 12 | 13 | 0 | 0 | 25 | XP_001446881 |
| PTET.51.1.P1180147 | 7 | 16 | 0 | 0 | 23 | XP_001427491 |
| PTET.51.1.P006001 | 9 | 10 | 3 | 0 | 22 | XP_001452484 |
| PTET.51.1.P0620215 | 10 | 11 | 0 | 0 | 21 | XP_001453643 |
| PTET.51.1.P0170077 | 11 | 9 | 0 | 0 | 20 | XP_001449252 |
| PTET.51.1.P0400053 | 12 | 8 | 0 | 0 | 20 | XP_001436180 |
| PTET.51.1.P0450314 | 6 | 12 | 0 | 0 | 18 | XP_001448195 |
| PTET.51.1.P0080258 | 8 | 10 | 0 | 0 | 18 | XP_001458158 |

